# RNAi-based discrimination of exogenous DNA by mitotic heterochromatin in fission yeast

**DOI:** 10.64898/2026.03.26.714489

**Authors:** Hirohisa Ebina, Mattia Valentini, Haochen Yu, Lisa Baumgartner, Karl C.F. Olsen, Rajendran Rajeswaran, Sigurd Braun, Olivier Voinnet, Yves Barral

## Abstract

Various mechanisms protect prokaryotic cells from invasive exogenous (exo)DNA. However, these are rarely conserved in eukaryotes, where the existence of such processes remains elusive. Here, we uncover how fission yeast eliminates exoDNA through unequal partitioning. We show that intrinsic transcriptional features of plasmidial exoDNA produce small interfering (si)RNAs that recruit heterochromatin onto the plasmid. An active, heterochromatin-dependent clustering mechanism then causes its asymmetric partitioning during mitosis. The mitotic kinase Aurora B, which dissociates heterochromatin on segregating chromosomes, is distinctively hypoactive on plasmidial DNA, facilitating its unequal partitioning and, ultimately, elimination from the cell population. Thus, an interplay between RNAi-mediated exoDNA heterochromatinization and chromosomal phosphorylation enables fission yeast to discriminate self-from non-self-DNA, uncovering a hitherto undescribed cell-autonomous immune process in eukaryotic cells.

## Main Text

Various cell-autonomous mechanisms, including the restriction-modification and CRISPR-Cas systems (*1, 2*), protect prokaryotes from invasive exogenous (exo)DNA. In contrast, the fate of exoDNA is comparatively much less well-characterized in eukaryotic cells. In some animal cells, transfected DNA is rapidly silenced and eliminated upon its clustering and encapsulation in a membrane-bound cytoplasmic “exclusome” (*3*). In budding yeast (*Saccharomyces cerevisiae*), exoDNA remains confined within mother cells during mitosis, causing its rapid elimination from the cell population, unless it contains a dedicated ∼120 nt *CEN* sequence (*4–7*). While the exclusome-based mechanisms in metazoans and the confinement-based mechanisms in *S. cerevisiae* are being progressively elucidated (*3, 8–12*), understanding the broader principles of exoDNA discrimination and elimination in eukaryotes requires investigating how other species accomplish this. The symmetrically dividing cells of fission yeast (*Schizosaccharomyces pombe*), for instance, rapidly eliminate plasmid exoDNA, even if replicative, albeit via unknown mechanisms (*13–15*).

Protective cellular mechanisms also exist against invasive exoRNA. These include RNA interference (RNAi) mediated by small interfering RNAs (siRNAs) processed from double-stranded precursors (dsRNAs). Virus-derived siRNAs confer antiviral immunity (*16*), while transposon-derived siRNAs help maintain genome integrity by guiding heterochromatin formation at their loci of origin (*17*). In the transposon-poor *S. pombe* genome (*18*), RNAi seems repurposed for heterochromatin assembly at pericentromeric repeats (*19*). This is required for centromere function because RNAi-deficient *S. pombe* mutants exhibit aberrant chromosome segregation (*20*). However, protective mechanisms targeting exoRNA or exoDNA have not yet been identified in fission yeast. Here, we investigated whether RNAi or any dedicated cellular mechanism mediates the discrimination and elimination of exoDNA in fission yeast.

### Mitotic *S. pombe* cells actively partition exoDNA asymmetrically via clustering at the nuclear periphery

We used the replicative *TetO* repeat-containing plasmid pYB1761 to monitor the fate of exoDNA in *S. pombe* (Fig.1A). Transformed into cells expressing the GFP/mCherry-fused TetO-binding protein TetR (TetR-GFP/mCherry), pYB1761 formed mostly nuclear dots under fluorescence microscopy (Fig.1B). The dots varied in number between zero and >20 and had variable intensities likely reflecting varying plasmid copy numbers therein (Fig.1C-D). In most late mitotic cells, one daughter cell often contained numerous and intense dots, while the other often lacked detectable signals entirely (Fig.1B). We thus set out to explore whether this apparent unequal partitioning involved passive, random partitioning or active mechanisms. Image-based sorting during late mitosis delineated three major occurrences: (i) fully asymmetric partitioning, where one daughter receives all dots; (ii) partially asymmetric partitioning, where one daughter receives more than twice as many dots as the other; or (iii) (near) symmetric partitioning (Fig.1E). Strikingly, full asymmetry (i) was 71% more frequent in populations than expected from random segregation (Fig.1F) *i.e*. greater than four standard deviations above the theoretical expectation (Fig.1G) with polynomial distribution as the null hypothesis (*Z* = 4.11, *p* < 10^−4^; methods).

**Fig. 1.**
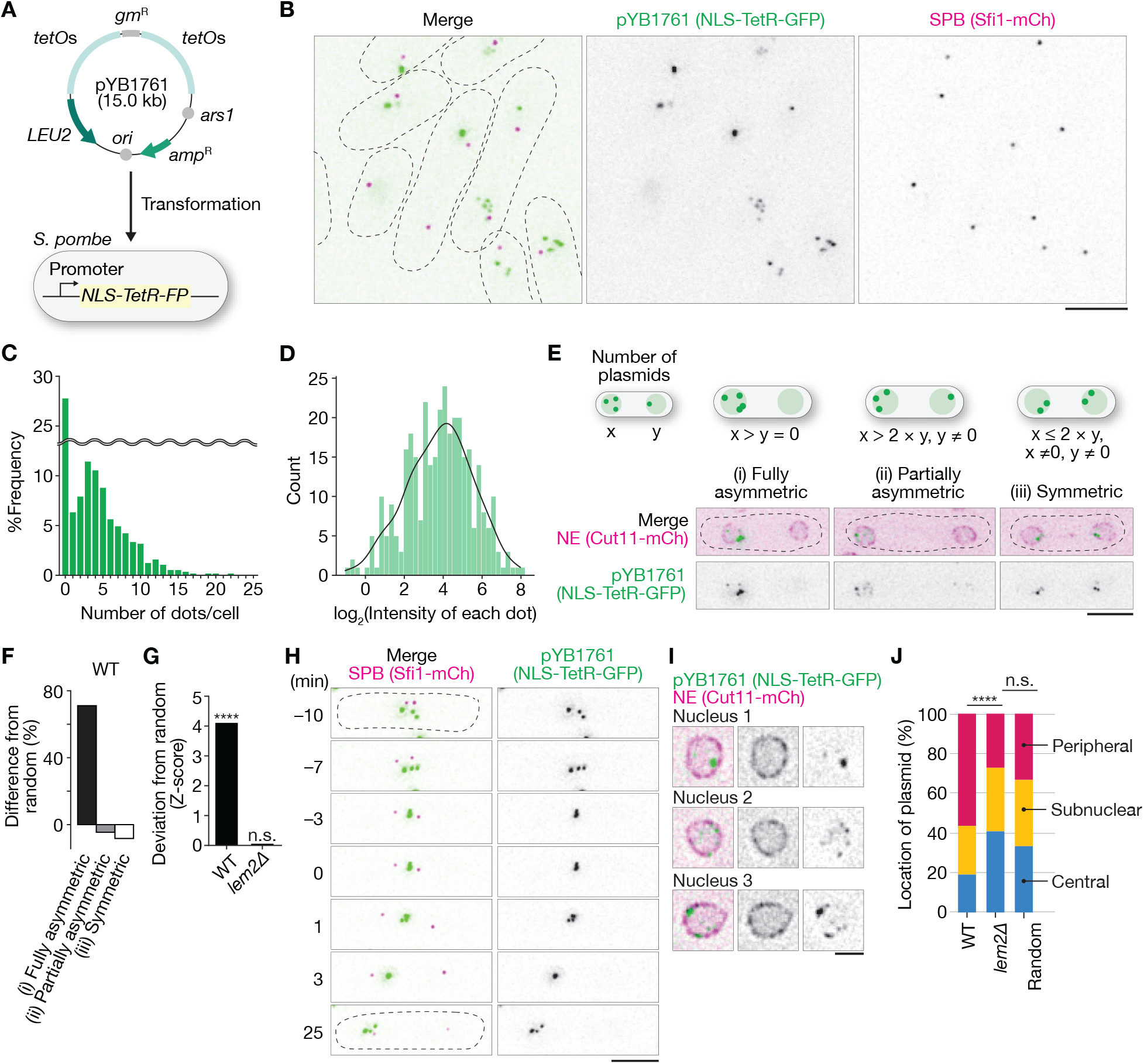
Fission yeast cells partition the plasmid asymmetrically during mitosis. (**A**) Cartoon of a replicative plasmid (pYB1761) used in this study. pYB1761 carries an array of 182 *tetO* repeats in total, a replication origin active in fission yeast (*ars1*), an auxotrophic selection marker taken from budding yeast (*LEU2*), two bacterial selection markers (*amp*^R^ and *gm*^R^), and a replication origin active in bacteria (*ori*). (**B**) Microscopy images of wild-type cells showing pYB1761 (green) with SPB (magenta). The dotted lines represent the cell outlines. Scale bars, 5 µm. (**C**) Quantification of the frequency of cells with each number of dots in wild-type cells. *n* = 569 cells. (**D**) Quantification of the intensity of each dot in wild-type cells. A kernel density estimate of the histogram was drawn as a black line. *n* = 338 dots. (**E**) Microscopy images of each partition pattern of pYB1761 are shown, along with schematic illustrations above. The definition of each partition pattern was provided in the form of equations. Scale bars, 5 µm. (**F**) Percentage of difference in the fraction of each partition pattern of pYB1761 compared to the theoretically expected fraction calculated from the random partitioning in wild-type cells. *n* = 1251 cells. For details, see Materials and Methods. (**G**) Quantification of the divergence of the distribution of pYB1761 partition patterns from the expected as a function of the theoretical standard deviation (*Z*-score) in wild-type cells. *n* > 492 cells. ^****^*p* < 0.0001 (binomial test analyses). For details, see Materials and Methods. (**H**) Microscopy images of a representative wild-type cell showing the fully asymmetric partitioning of pYB1761 (green) with SPB (magenta). The time point 0 min corresponds to the onset of anaphase B, where two SPBs restart moving towards each cell end. Scale bars, 5 µm. (**I**) Microscopy images of pYB1761 (green) localizing to the periphery of the nuclear envelope (NE; magenta) from three different nuclei. Single sections of the deconvoluted image were shown. Scale bar, 2 µm. (**J**) Quantification of pYB1761 distribution relative to the nuclear periphery. The nuclear area of each section of the microscopy images was divided into three zones with equal area. *n* > 655 dots. n.s., not significant; ^****^*p* < 0.0001 (χ^2^ test analyses).

In live-cell imaging conducted over several divisions, asymmetric plasmid partitioning was independent of the orientation of the old versus new spindle pole bodies (SPB), as assessed with Sfi1-CFP and Cdc7-GFP (Fig.S1A-C) (*21–23*), and also independent of the old and new cell ends (Fig.S1D-E). The plasmid’s presence did not alter mitosis progression, save a slight increase in metaphase spindle length (Fig.S1F-I), making mitotic defects an unlikely cause of asymmetry. In a time-course analysis (Fig.1H), plasmids tended to dynamically congregate into one or a few brighter fluorescent dots. Mitosis exacerbated this phenomenon, causing clustered plasmids to partition more in one given daughter cell during late anaphase. Clusters then tended to dissociate into individual fluorescent foci at, or shortly after, mitotic exit (Fig.1H), coinciding with the DNA replication onset. Plasmid clusters remained nuclear during the entire mitotic process, which does not involve nuclear envelope breakdown in *S. pombe* (*24*). They also localized preferentially to the nuclear periphery (Fig.1I-J) in a manner requiring the inner nuclear membrane protein Lem2/Heh1 (Lap-Emerin-Man domain proteins2/Helix-Extension-Helix domain protein 1) (*25*– *30*). We conclude that a hitherto unknown active mechanism that likely causes their physical clustering at the nuclear periphery governs unequal partitioning of exoDNA plasmid copies in dividing *S. pombe* cells.

### Histone H3 Lysine 9 is methylated on a discrete exoDNA plasmid’s region

Lem2 organizes heterochromatin at the nuclear periphery (*25, 26, 30–32*). Moreover, heterochromatin self-cohesion promotes coalescence of heterochromatinized genomic regions into clusters (*33–35*). Thus, we asked if plasmid heterochromatinization causes plasmid clustering. Methylation of histone H3 lysine 9 (H3K9me) by Clr4, the methyltransferase Suv39 homolog in *S. pombe*, marks the initiation of heterochromatin assembly, which occurs at telomeres, pericentromeric repeats, and the silent mating type locus (Fig.2A). There, an Ago1-associated RNAi effector complex, loaded with repeat-derived siRNAs generated by the Dicer ribonuclease (Dcr1), guides Clr4 recruitment. Subsequent binding of Swi6/HP1α, the main eukaryotic heterochromatin scaffold, to H3K9me ensures heterochromatin coalescence and compaction (Fig.2A) (*36–41*).

**Fig. 2.**
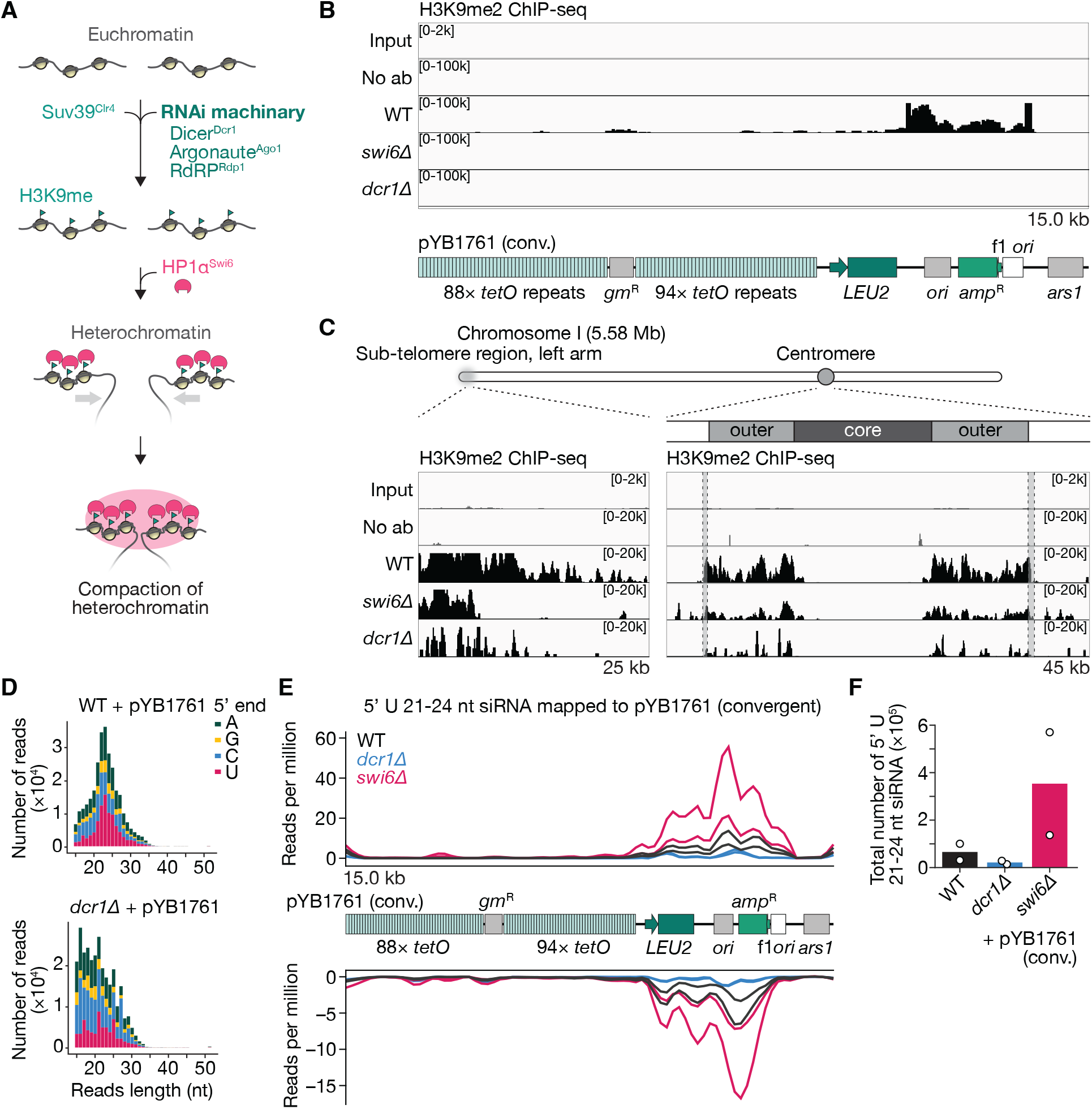
RNAi-dependent heterochromatin formation on the plasmid. (**A**) Cartoon of a simplified mechanism of RNAi-dependent heterochromatin formation on chromosomes in *S. pombe*. (**B**) Profiles on H3K9me2 ChIP-seq on pYB1761 for the indicated genotypes. No ab, negative control without antibodies. The linearised map of pYB1761 is depicted at the bottom. (**C**) Profiles of H3K9me2 ChIP-seq on the sub-telomeric region on the left arm of chromosome 1 (left) and the centromeric region of chromosome 1 (*cen1*, right). The dotted squares with grey colour indicate the inverted repeat elements that flank *cen1* (IRC1). No ab, negative control without antibodies. (**D**) 5’-terminal nucleotide composition of deep-sequenced sRNAs from wild-type and *dcr1Δ* cells, which were transformed with pYB1761. (**E**) Profile of 5’U 21-24 nt siRNA from the indicated genotype mapped to pYB1761. *N* = 2 trials for each of the indicated genotypes. (**F**) Plot of the total number of 5’U 21-24 nt siRNA mapped to pYB1761 in each genotype. *N* = 2 trials for each of the indicated genotypes.

Chromatin immunoprecipitation followed by sequencing (ChIP-seq) using an H3K9me2-specific antibody indeed revealed strong plasmidial H3K9me2 enrichment, yet almost exclusively between the *LEU2* and *ars1* sequences, where it substantially overlapped with a region flanked by the P.*LEU2* and P.*amp*^R^ convergent promoters (Fig.1A;2B). As expected, H3K9me2 was also detected on chromosomes at the pericentromeric and sub-telomeric but not euchromatic regions (Fig.2C;S2). Both plasmidial and chromosomal H3K9me2 signals dropped in *dcr1Δ* and *swi6Δ* mutant cells. We conclude that a discrete region of the plasmid is non-randomly labelled for heterochromatin assembly in an RNAi- and Swi6-dependent manner.

### Dicer-dependent siRNA accumulation specifically overlaps the plasmidial H3K9me2 signal

To test whether plasmid-mapping siRNAs accumulate in pYB1761-transformed cells and overlap the H3K9me2-enriched region, we deep-sequenced total small (s)RNAs extracted from pYB1761-transformed wild-type, *dcr1Δ*, or *swi6Δ* cells (Fig.S3A). More than 0.3% of total sRNA reads mapped to the plasmid as opposed to the genome, a substantial fraction given the small plasmid size (∼0.1%). Plasmid-mapping sRNAs had a discrete 20-27-nt size-distribution peaking at 22-23 nt (Fig.2D;S4), typical of *S. pombe* Dcr1 products (*42, 43*). These sRNAs specifically mapped onto the H3K9me2-enriched region flanked by P.*LEU2* and P.*amp*^R^ (Fig.S3B-C), though their distribution extended slightly further into the *LEU2* coding sequence. Their mapping onto both DNA strands was consistent with dsRNA origins. Selective mapping of 21-24 nt sRNAs with a 5’ uridine (hereafter siRNAs), which are preferentially bound by *S. pombe* Ago1 (*39, 43*), yielded a near-identical profile (Fig.2E;S3B). Moreover, the size-distribution pattern and abundance of plasmid-mapping siRNAs were substantially altered and strongly reduced respectively in *dcr1*Δ mutant cells carrying pYB1761 (Fig.2D-F;S3B-C;S4). As expected, pericentromeric repeat-derived siRNAs (*44*) were essentially eliminated in *dcr1*Δ cells (Fig.S3D). Therefore, bona fide Dcr1-dependent siRNAs specifically match the H3K9me2-enriched plasmid region. Also as expected, pericentromeric siRNAs vanished in pYB1761-transformed *swi6*Δ mutant cells (Fig.S3D) (*38, 45, 46*). Unexpectedly, however, the levels of plasmid-mapping siRNAs strongly increased in these cells (Fig.2E-F;S3B-C;S4), suggesting that Swi6-dependent silencing of plasmid transcription limits siRNA accumulation over the H3K9me2-enriched region. Collectively, these results support a model whereby siRNA-mediated heterochromatin assembly of plasmid exoDNA enables its clustering and unequal partitioning during mitosis. They also suggest that plasmid transcription is key to this process and is limited by it.

### RNAi- and heterochromatin-deficient cells show defects in plasmid clustering, asymmetric partitioning, and population-level elimination

The above observations predicted that *S. pombe* cells deficient in RNAi and heterochromatin formation would fail to cluster the plasmid and, hence, fail to partition it asymmetrically. Compared to wild-type cells, *swi6Δ* and *clr4Δ* mutants transformed with pYB1761 indeed showed more plasmid dots with reduced fluorescence intensity (Fig.3A-C). The *dcr1Δ* and *ago1Δ* mutant cells showed similar trends (Fig.S5). Moreover, the partitioning of the dots became indistinguishable from random in either of the *swi6Δ* (*Z* = 0.08, *p* = 0.52), *clr4Δ* (*Z* = 0.79, *p* = 0.26), *dcr1*Δ (*Z* = 0.39, *p* = 0.40), and *ago1Δ* (*Z* = 0.26, *p* = 0.44) mutant cells (Fig.3D).

**Fig. 3.**
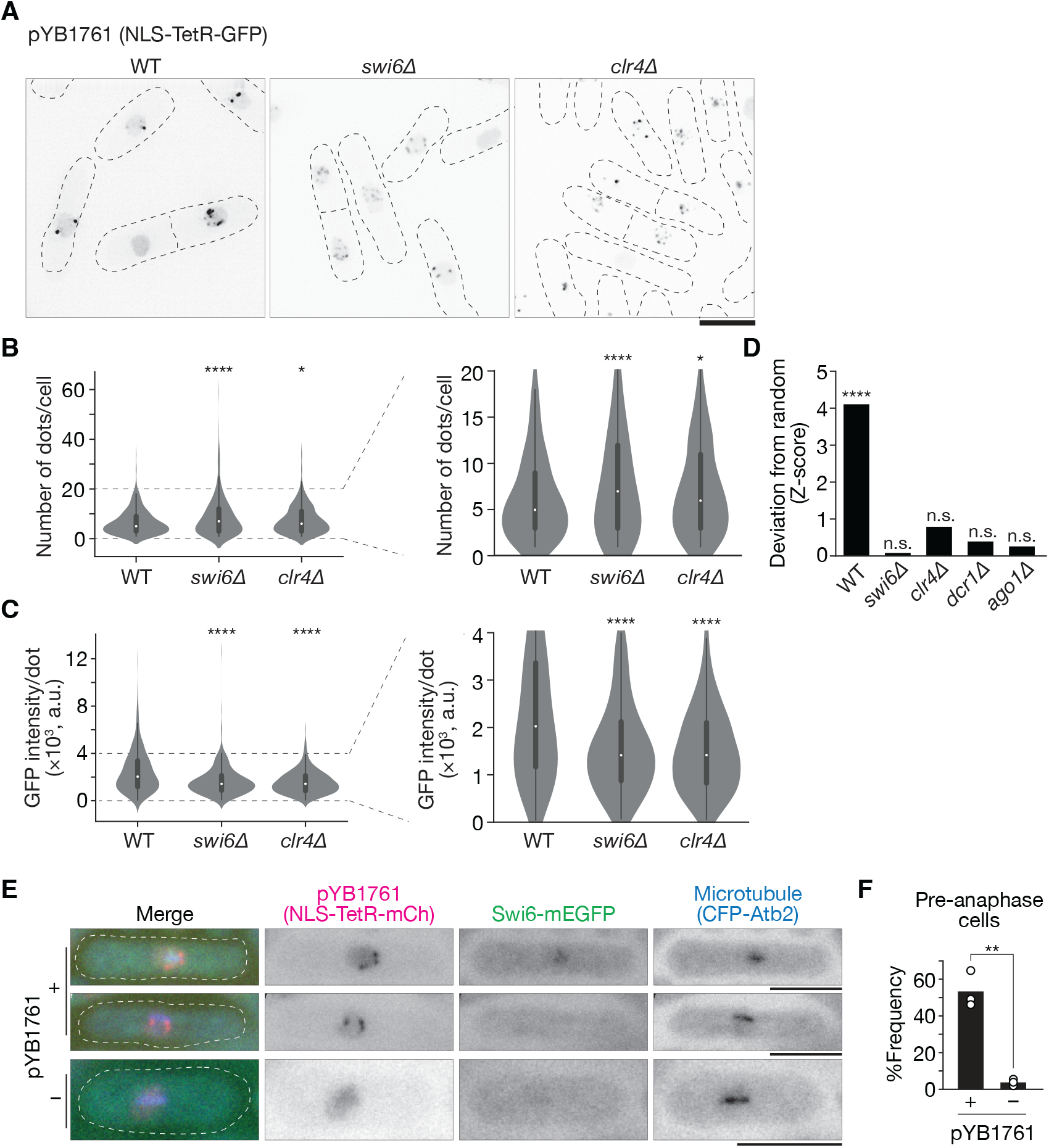
RNAi-dependent heterochromatin machinery is required for clustering and asymmetric partitioning of the plasmid. (**A**) Microscopy images of pYB1761 in the indicated genotypes. The black dotted lines represent an outline of cells. (**B**) Left, quantification of the number of fluorescent dots in the indicated genotypes. Right, same as the left, except zoomed in on the indicated range. *n* > 392 cells were tested. A white circle in each violin plot represents the mean value. ^*^*p* < 0.05; ^****^*p* < 0.0001 (Welch’s *t*-test). (**C**) Left, quantification of fluorescent intensity of each dot in the indicated genotypes. Right, same as the left, except zoomed in on the indicated range. *n* > 321 dots were tested. A white circle in each violin plot represents the mean value. ^****^*p* < 0.0001 (Welch’s *t*-test). (**D**) Quantification of the divergence of the distribution of pYB1761 partition patterns from the expected in indicated cells. Data for wild-type cells is the same as Fig. 1G. *n* > 209 cells. n.s. not significant; ^****^*p* < 0.0001 (binomial test analyses). For details, see Materials and Methods. (**E**) Microscopy images of pre-anaphase cells showing ectopically-expressed Swi6-mEGFP (green), and microtubules (blue) with or without pYB1761 (magenta). (**F**) Quantification of the frequency of pre-anaphase cells showing Swi6-GFP dots. *n* > 101 cells were tested in each trial. *N* = 3 trials. ^**^*p* < 0.01 (Student’s *t*-test). Scale bars, 5 µm.

Given Swi6 functions as a heterochromatin scaffold protein, the model predicts its association with pYB1761 in mitotic cells. Cells expressing an extra-copy of Swi6 C-terminally fused to mEGFP (Swi6-mEGFP) were therefore monitored as they entered mitosis and assembled a metaphase spindle, visualized by CFP-labelled α-tubulin (Fig.S1G). In these mitotic cells, which partition plasmids asymmetrically similar to wild-type cells (Fig.S6), Swi6-mEGFP-decorated structures formed in cells containing the plasmid, detected via TetR-mCherry, but were absent in cells that had lost it (Fig.3E-F). The Swi6-mEGFP decoration remained adjacent to pYB1761 dots but did not directly colocalize with them, possibly reflecting that the large plasmidial *tetO* array is not itself siRNA-targeted and heterochromatinized (Fig.2B;2E).

To explore the impact of RNAi and heterochromatin on exoDNA elimination at the population level, cells growing exponentially under non-selective conditions (*i.e*., in the presence of leucine, yielding a division time of ∼3hrs) were imaged every 6 hours after release from the selection (Fig.S7A-B). The proportion of plasmid-containing cells was then quantified using the TetR-GFP signal. Both wild-type and mutant cell populations eventually lost pYB1761 over time, yet this was substantially slower in *swi6Δ* and *dcr1*Δ mutants than in the wild-type strain (Fig.S7C) despite their comparable growth rates (Fig.S7D-E). Collectively, these results suggest that siRNA-guided heterochromatinization enables plasmid copies to coalesce into heterochromatic clusters. This likely underpins their unequal partitioning during mitosis, ultimately allowing fission yeast to eliminate exoDNA at the population level.

### Plasmid-intrinsic transcriptional features govern the extent of siRNA production and asymmetric plasmid partitioning

Both H3K9me2- and siRNA-signals coincidently mapped onto the region delineated by the P.*LEU2* and P.*amp*^R^ convergent promoters (P.*LEU2*→←P.*amp*^R^). *S. pombe* Dcr1 can process siRNAs from dsRNA produced by sense/antisense genomic transcription (*47*). Thus, P.*LEU2*→←P.*amp*^R^ might constitute a key plasmid-intrinsic feature for dsRNA production, siRNA processing, heterochromatinization, and asymmetric partitioning. Alternatively, asymmetric plasmid partition may indirectly result from the heterochromatinization of chromosomal, *i.e*. plasmid-extrinsic rather than the plasmidial locus. To distinguish between these possibilities, we engineered a variant of the pYB1761 plasmid, pYB2381, in which the orientation of the *LEU2* gene is reversed (P.*LEU2*←←P.*amp*^R^) to prevent convergent transcription, while leaving the remainder of the original plasmid unchanged (Fig.4A). Deep-sequencing detected 5’U-terminal 21-to-24-nt plasmid-mapping siRNAs exclusively onto the modified P.*LEU2*←←P.*amp*^R^ region in pYB2381-transformed wild-type cells (Fig.4B-C;S3E-F;S4). However, their levels were far below those of siRNAs mapping the P.*LEU2*→←P.*amp*^R^ region in pYB1761-transformed wild-type cells. Thus, while convergent transcription likely provides the bulk of plasmid-derived siRNAs, an siRNA source exists independently of it. In pericentromeric repeat silencing, RNA-dependent RNA polymerase (RdRP; Rdp1 in *S. pombe*) produces dsRNA *de novo* from repeat-derived single-stranded transcripts (*45, 48*) (Fig.S3D), a mechanism possibly at work on the P.*LEU2*←←P.*amp*^R^ region in pYB2381. Rdp1 also likely produces low dsRNA levels in pYB1761-transformed wild-type cells because plasmid-mapping sRNA accumulation was slightly but consistently reduced in pYB1761-transformed *rdp1*Δ mutant cells (Fig.4D-E;S3G-H;S4). We conclude that plasmid-derived dsRNA production depends on plasmid-intrinsic transcriptional features, including convergent transcription and, less prominently, Rdp1 activity.

**Fig. 4.**
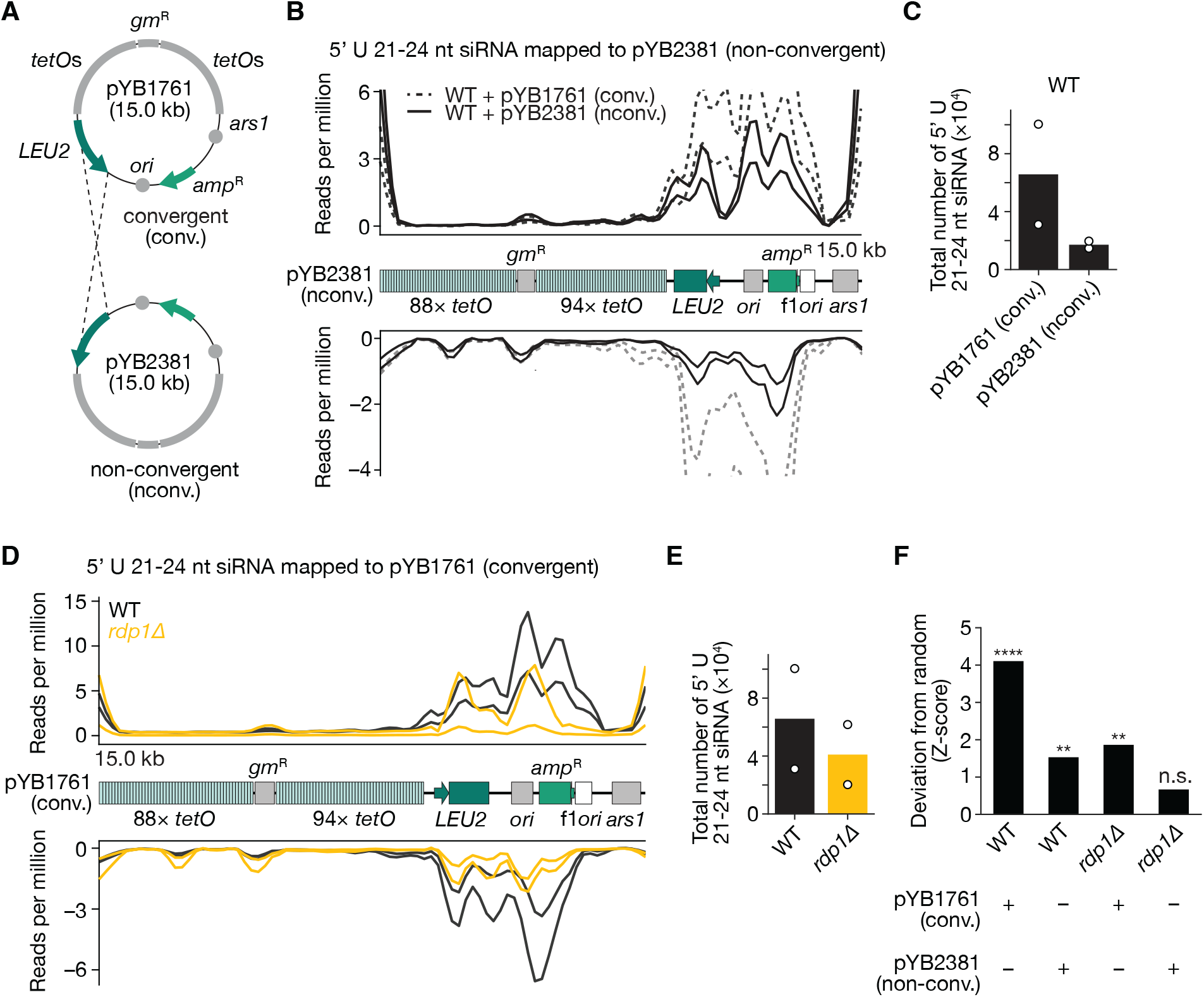
Impacts of plasmid-intrinsic transcription features on sRNA production and asymmetric partitioning of the plasmid. (**A**) Cartoon of the convergent plasmid (pYB1761) and a new non-convergent plasmid (pYB2381). (**B**) Profile of 5’U 21-24 nt siRNA from wild-type cells mapped to pYB1761 (dotted lines) or pYB2381 (solid lines). *N* = 2 trials for each condition. (**C**) Plot of the total number of 5’U 21-24 nt siRNA mapped to pYB1761 or pYB2381 in wild-type cells. *N* = 2 trials for each of the indicated genotypes. (**D**) Profile of 5’U 21-24 nt siRNA from the indicated genotype mapped to pYB1761. *N* = 2 trials for each of the indicated genotypes. (**E**) Plot of the total number of 5’U 21-24 nt siRNA mapped to pYB1761 in the indicated genotypes. *N* = 2 trials for each of the indicated genotypes. (**F**) Quantification of the divergence of the distribution of pYB1761 partition patterns from the expected in indicated cells. Data for wild-type cells with pYB1761 is the same as Fig. 1G. *n* > 518 cells. n.s. not significant; ^**^*p* < 0.01; ^****^*p* < 0.0001 (binomial test analyses). For details, see Materials and Methods.

If plasmidial rather than chromosomal loci drive clustering and unequal partitioning, then preventing convergent transcription, which strongly reduces siRNA levels (Fig.4B-C), would be expected to compromise heterochromatin assembly of the plasmid and therefore randomize its mitotic distribution. Supporting this idea, H3K9me2 levels were reduced, but not abolished, in the P.*LEU2*←←P.*amp*^R^ region of pYB2381 compared to the P.*LEU2*→←P.*amp*^R^ region of pYB1761 (Fig.S8). Furthermore, pYB2381 partitioned significantly less asymmetrically than pYB1761 (*Z* = 2.93, *p* < 10^-2^ vs *Z* = 4.11, *p* < 10^−4^, Fig.4F). Partitioning of pYB2381 was fully randomized in the *rdp1*Δ mutant cells (Fig.4F), consistent with Rdp1’s involvement in pYB2381 asymmetric partitioning, as hypothesized above. Therefore, the extent of unequal plasmid partitioning correlates with plasmid-derived siRNA levels. We conclude that plasmid-intrinsic, RNAi-dependent heterochromatin assembly and clustering are the mechanisms underlying asymmetrically plasmid partitioning during mitosis in *S. pombe*.

### Aurora B/Ark1 discriminates chromosomal from exoDNA heterochromatin

The above findings were intriguing because heterochromatin does not impede symmetric segregation of chromosomes. Indeed, chromosomal heterochromatin, particularly on centromeres, is dissolved during mitosis owing to phosphorylation of H3-Serine10 neighbouring the H3K9me2 mark by the Aurora B kinase (Ark1 in *S. pombe*) (*49, 50*). To explore this apparent plasmid-versus-chromosome discrepancy, we used antibodies raised against histone H3 or S10-phosphorylated H3 (H3S10p) in pYB1761-transformed, metaphase wild-type cells. In quantitative PCR-based ChIP (ChIP-qPCR) assays corrected for the plasmid’s reduced histone abundance, the plasmid H3K9me2-enriched region had much less H3S10p signal than chromosomal heterochromatic or even euchromatic regions (Fig.5A). Thus, Ark1 barely phosphorylates the plasmid-associated histone H3S10 during mitosis. Furthermore, targeting Ark1 to the plasmid using a TetR-fusion (Ark1-TetR;Fig.5B-C;S9) randomized mitotic plasmid partitioning, unlike overexpression of unfused Ark1. By contrast, a catalytic-dead Ark1-TetR allele (Ark1-K118R-TetR) (*51*) did not promote mitotic randomization (Fig.5C). Therefore, unequal partitioning in wild-type cells relies on plasmid-associated H3S10 evading phosphorylation by Ark1, further strengthening the notion that the plasmid-intrinsic heterochromatin, but not heterochromatin elsewhere in the genome, promotes its unequal partitioning. We conclude that Ark1 acts as a discriminator of endogenous versus exogenous heterochromatic DNA.

**Fig. 5.**
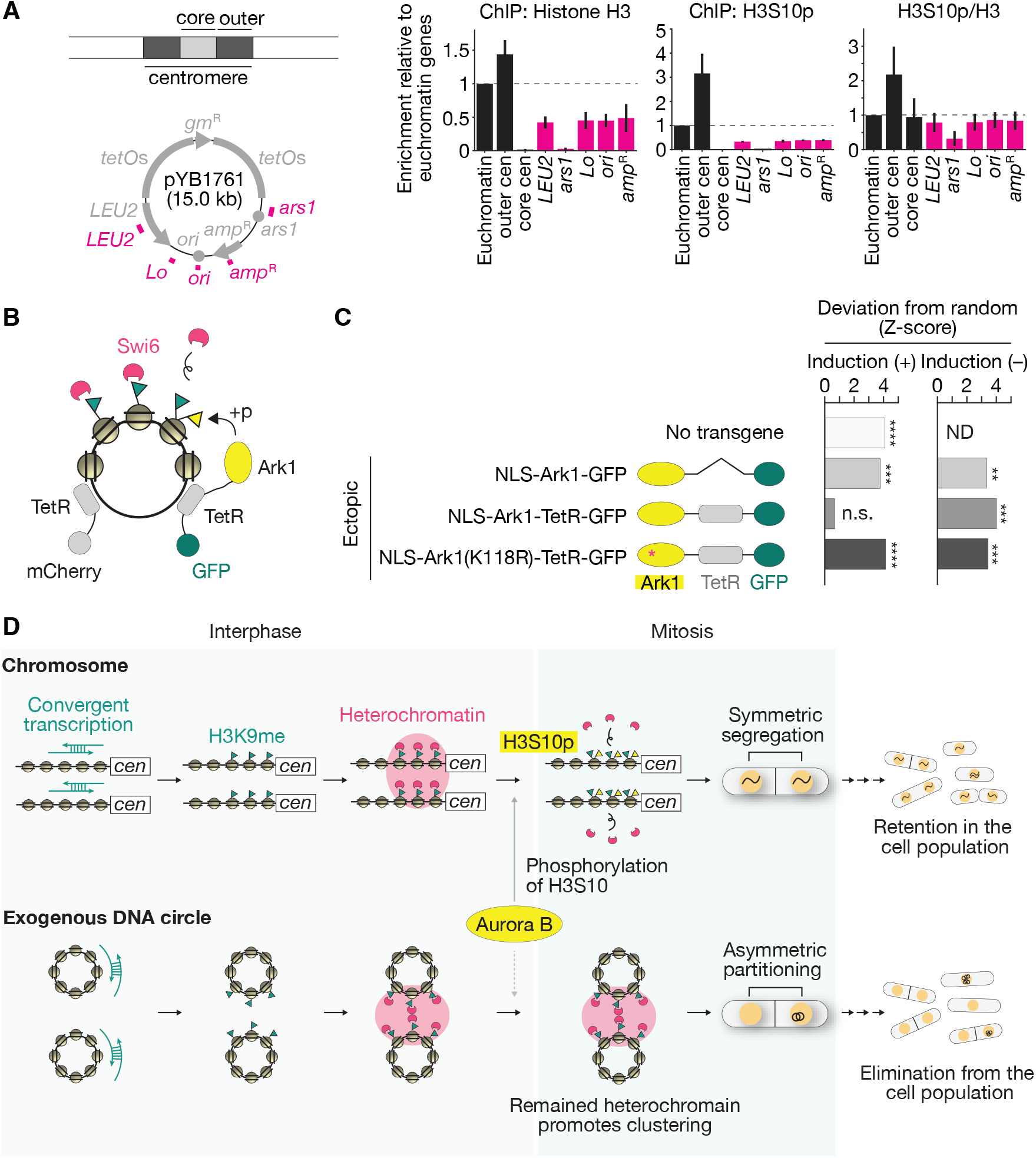
Aurora B does not phosphorylate the plasmid. (**A**) Histone H3 and H3S10p on indicated regions of chromosomes and pYB1761 (shown as magenta lines in the cartoon) were detected by ChIP-qPCR in metaphase wild-type cells. Error bars, SEM of *N* = 3 experiments. (**B**) Cartoon of Ark1 tethering on pYB1761. (**C**) Quantification of the divergence of the distribution of pYB1761 partition patterns from the expected in indicated cells. Data for wild-type cells without the transgene (No transgene) in the induced condition (+) is the same as Fig. 1G. *n* > 185 cells. n.s. not significant; ^**^*p* < 0.01; ^**^*p* < 0.001; ^****^*p* < 0.0001 (binomial test analyses). For details, see Materials and Methods. ND, not determined. (**D**) Model of RNAi-based discrimination and population-level elimination of exoDNA by mitotic heterochromatin.

## Discussion

Here, we have identified a hitherto unknown function for RNAi-guided heterochromatin assembly in *S. pombe:* the discrimination and elimination of exoDNA via asymmetric partitioning during mitosis. This is achieved through heterochromatin-mediated exoDNA clustering into discrete clusters, *de facto* increasing the probability that one of the two daughter cells inherits all or the greater portion of exoDNA. Reiterated over cell divisions, the process eventually eliminates exoDNA from the population (Fig.5D;S7). Plasmid-derived RNA is the primary determinant of this immunity because it is the source of siRNAs initiating heterochromatinization. Plasmid transcription is therefore essential and, indeed, Swi6 limits plasmid-derived siRNA accumulation presumably through transcriptional gene silencing of the plasmidial P.*LEU2*→←P.*amp*^R^ unit in pYB1761 (Fig.2E-F;S4). On chromosomal pericentromeric repeats, by contrast, Swi6 initiates and presumably maintains siRNA production by indirectly recruiting Rdp1 subunit Hrr1 (*52, 53*). Therefore, while Swi6 is required for both plasmidial and chromosomal heterochromatin assembly, its respective positive and negative roles on siRNA production represent a discriminator of exo-versus endo-DNA that deserves further investigation.

Decreased siRNA levels from the non-convergent pYB2381 plasmid correlated with lower asymmetric partitioning, indicating an siRNA-dose-dependent mechanism. Unequal pYB2381 partitioning was abrogated in *rdp1*Δ mutant cells, suggesting that the Rdp1-dependent conversion of plasmid-derived single-stranded (ss)RNA into dsRNA can initiate the process. Unlike pYB1761, plasmidial exoDNA would not routinely contain converging transcription units. Therefore, Rdp1 might be a key and generic component of *S. pombe’s* immunity against exoDNA, raising the question of how Rdp1 activity might be stimulated. Remarkably, pYB2381-derived siRNAs mapped precisely to the P.*LEU2*←←P.*amp*^R^ region and were qualitatively similar to the pYB1761-derived species (Fig.4B). Rdp1 also contributed to siRNA biogenesis from P.*LEU2*→←P.*amp*^R^ in pYB1761 (Fig.4D,E), suggesting that its recruitment is intrinsic to this region. Notably, the *amp*^R^ open reading frame therein is of bacterial origin. This might predispose its transcription into spurious polyA^-^ or uncapped mRNAs in yeast cells. Legit transcripts from *amp*^*R*^ might also display suboptimal codons, leading to ribosome stalling and ensuing mRNA breakage. Aberrant RNAs resulting from either scenario are known RdRP substrates in various organisms (*54–56*). Abnormal nucleosome arrays might also form over *amp*^*R*^ and enforce Rdp1 recruitment thereupon via as-yet-undetermined mechanisms. It is therefore interesting that the other bacterial *gm*^R^ open reading frame embedded in the *tetO* arrays did not generate siRNA and likely no Rdp1 targeting (Fig.4B,D). What differentiates these two bacterial genes remains elusive.

Restricting Aurora B localisation and its activity to chromosomes emerges as a complementary means of distinguishing endo-from exoDNA, ultimately ensuring that only the former is unequally partitioned during mitosis. Therefore, self-versus-non-self-discrimination in fission yeast is conceptually reminiscent of that of prokaryotic restriction-modification systems, where specific modification of the self also protects it from elimination. Strikingly, the immunity function of Aurora B is conserved in evolutionarily distant fungi (*10*), suggesting, perhaps, an ancestral role for Aurora B in self-versus-non-self-DNA discrimination. Together, our data suggest DNA-immunity mechanisms in eukaryotes might have co-evolved with other key eukaryotic features, such as mRNA processing, RNAi, chromatin structure, and the mitotic machinery, where these mechanisms are now hidden in plain sight.

## Acknowledgments

We are grateful to A. Stevenson and T. C.T. Michaels for developing and discussing the statistical analyses used in this work, P. Leary from FGCZ for his help with sRNA-seq bioinformatic analyses, and H. Tsuyuzaki and M. Sato for providing us with the plasmid pECBZ2-bsd used for integrating Ark1-tethering constructs in the *z2* locus. H.E. acknowledges D. Qiu for assistance with constructing Ark1-tethering constructs. We also thank S. Martin for help during the earlier steps of the project. We acknowledge the ScopeM facility at the Institute of Biochemistry, ETH Zürich, for their technical support and assistance in this work. We would like to thank all members of the Barral and Kroschewski lab for their valuable comments on the work and the manuscript.

## Author contributions

Conceptualization: H.E., M.V., H.Y., O.V., and Y.B.

Formal analysis: H.E., M.V., K.C.F.O., R.R.

Investigation: H.E., M.V., H.Y., K.C.F.O., R.R., with contributions from L.B. Methodology: H.E., M.V., H.Y.

Supervision: Y.B. Visualization: H.E., M.V.

Writing – original draft: H.E., Y.B.

Writing – review & editing: H.E., K.C.F.O., R.R., S.B., O.V., Y.B.

## Data and materials availability

The ChIP-seq data for this study have been deposited in the European Nucleotide Archive (ENA) at EMBL-EBI under accession number PRJEB97961. The small RNA-seq data for this study have been deposited in the Sequence Read Archive at NIH under accession number PRJNA1334196. All unique materials and reagents generated in this study are available upon request.

## Supplementary Materials

Materials and Methods

Figs. S1 to S9

Tables S1 to S3

