## Supplemental information for "RNAi-based discrimination of exogenous DNA by mitotic heterochromatin in fission yeast"

### Materials and Methods

#### Yeast strains, media, and genetics

The strains and plasmids used in this study are listed in Tables S1 and S2, respectively. Nomenclatures are based on the previous paper (*57*). Standard methods for yeast genetics and gene tagging were used (*58–60*). For constructing the convergent plasmid pYB1761 and the non-convergent plasmid pYB2381, the *tetO* repeat array was cloned from pLAU44 (*61*). For expressing NLS-TetR-GFP under the thiamine-repressive *nmt81* promoter (*62*), a plasmid pYB1521 was linearized by AvrII and integrated into the *ura4* locus. For expressing NLS-TetR-GFP under the *nup211* promoter, a plasmid pYB2125 was linearized by AvrII and integrated into the *ura4* locus. For expressing NLS-TetR-mCherry under the *nup211* promoter, a plasmid pYB2144 was linearized by AvrII and integrated into the *ura4* locus.

For ectopic expression of Swi6-mEGFP under the endogenous promoter of the *swi6*$^{+}$ gene (Fig.3E-F;S6), a plasmid pYB4098 was used for generating DNA fragments to replace the *SPAC25B8.09*$^{+}$ gene locus on chromosome I by transformation.

For ectopic expression of Ark1 constructs (Ark1-NLS-TetR-GFP, Ark1-TetR-GFP, or Ark1(K118R)-NLS-TetR-GFP) under the moderate conditions (Fig.5C;S9), a plasmid pYB4042, pYB4116, or pYB4084 was linearized by FseI and integrated into the *z2* locus on chromosome II, respectively (*63*, *64*). To avoid the unwanted effects of the ectopic Ark1 construct, the expression of the Ark1 constructs was suppressed by supplementing with 2 μg/mL of thiamine (Sigma). For inducing the Ark1 constructs, cells were pre-cultured overnight in Edinburgh minimal media 2 (hereafter EMM) containing leucine dropout and 2 μg/mL thiamine, and then washed twice with pre-warmed EMM–Leu. Then, cells were cultured in EMM–Leu medium without thiamine for 18-20 hours and subjected to microscopy (see below).

#### Microscopy

For fluorescent microscopy, yeast cells were cultured in EMM supplemented with five supplements (EMM+5S) or in Leucine dropout medium (EMM–Leu). 5-10 mL of cells from log-phase cultures were concentrated by centrifugation at 800 × *g*, and spotted on a round coverslip and immobilized with the EMM+5S (Fig.S7) or EMM–Leu agar patch. Live-cell imaging was performed using the DeltaVision-SoftWoRx system (Applied Precision) equipped with an Olympus IX71/81 inverse microscope, a 60× or 100×/1.4 NA objective, a CCD HQ2 camera

(Roper), 250 W Xenon lamps, and a temperature-controlled chamber. Images were acquired as serial sections along the z-axis, and deconvoluted and stacked using the "quick-projection" algorithm in SoftWoRx. The captured images were processed using Fiji (*65*).

For Fig.3E, 1 mL of cells from log-phase cultures was used for fixing with ice-cold 70% ethanol at –20°C for more than 6 hours and mounted on a microscope slide (Eperedia) for imaging.

**Synchronization of the cell cycle by hydroxyurea (HU)**

We transformed cells without expressing TetR with the indicated plasmid and picked up colonies from the selection plate (EMM–Leu) freshly. These colonies were pre-cultured at 30°C for 2 days and then inoculated on a large scale (200 or 300 mL) at 30°C overnight. Log-phase cultures were then diluted to OD595=0.4 and arrested by 12 mM of freshly prepared hydroxyurea (HU) (Sigma-Aldrich, #H8627) at 30°C for 4 hours with agitation (*66*). Arrested cells were washed twice with media preheated to 30 °C. OD595=0.4 cell cultures were prepared on the same scale and inoculated without HU at 30°C for 60 min for harvesting G2/M cells (sRNA-seq) or 90 min for harvesting metaphase cells (ChIP-qPCR, Fig.5A).

**Chromatin immunoprecipitation (ChIP) sequencing and ChIP-qPCR experiments**

Cells released from HU (for sRNA-seq and ChIP-qPCR in Fig.5A) or log-phase cells (for ChIP-seq and ChIP-qPCR in Fig.S8) were fixed with 1% (vol/vol) formaldehyde for 10 min with occasional agitation. Formaldehyde was quenched for 10 min by adding 125 mM glycine with occasional agitation. Cells were washed with ice-cold phosphate-buffered saline (PBS) twice, and then the cell pellets were quickly frozen in liquid nitrogen and stored at –80°C. Frozen cell pellets were resuspended in lysis buffer with proteinase and phosphatase inhibitors [50 mM HEPES/KOH (pH 7.5), 140 mM NaCl, 1 mM EDTA, 1% (vol/vol) Triton X-100, 0.1% (vol/vol) Na-deoxycholate, with inhibitors (1 mM AEBSF, 100 µg/mL Leupeptin, 4% of cOmplete™ protease inhibitor cocktail (Roche) suspended with PBS, 4% of PhosSTOP™ (Roche) suspended with PBS)] and were lysed by performing 7 cycles of 20 sec with the speed 6.0 m/sec followed by a 1 min cool-down on ice at 4°C using a FastPrep-24 5G (MP Biomedicals). Cell lysates were sonicated for 60 min by performing alternating cycles of 30-second pulses followed by a 30-second cool-down period at 4°C using a SONOPLUS HD 4050 Homogenizer (BANDELIN electronic). After multiple centrifugations for clarification, supernatants were immunoprecipitated at 4°C using anti-Histone H3 (Abcam, ab1791), anti-Histone H3 (phosphor S10) (Abcam, ab5176), and anti-H3K9me2 (Abcam, ab1220) antibodies and Dynabeads™ Protein G (Invitrogen, 10003D). We also included control IgG to account for non-specific binding in the ChIP fractions. Beads were first washed with lysis buffer without proteinase and phosphatase inhibitors twice, followed by washing with lysis buffer with 500 mM twice, then with wash buffer [10 mM Tris/HCl (pH 8), 250 mM LiCl, 1 mM EDTA, 0.5% (vol/vol) Nonidet P-40, 0.5% (vol/vol) Na-deoxycholate] twice, and then with Tris/EDTA (pH 8). Washed beads were suspended with elution buffer [50 mM Tris/HCl (pH 8), 10 mM EDTA, 0.8% (wt/vol) SDS]. Reverse crosslinking was performed by incubating first at 95°C at 1400 rpm for 10 min and then at 65°C at 500 rpm overnight. After proteinase K treatment, DNA fragments were purified using ChIP DNA Clean & Concentrator™ (Zymo Research, D5201). Input DNA samples were eluted with 50 µL of elution buffer, while ChIP DNA samples were eluted with 20 µL.

For ChIP-qPCR, targeted loci were amplified by qPCR using the Power SYBR Green PCR Master Mix (Applied Biosystems, A25742) and the StepOnePlus real-time PCR system (Applied Biosystems). Primers used for the qPCR experiments are described in Table S3. %Input was calculated in each target locus and each ChIP, then divided by the mean of three euchromatin loci (*act1*$^{+}$, *tef3*$^{+}$, and *ade2*$^{+}$) (*67*).

For ChIP-seq, once validated by ChIP-qPCR, libraries were constructed for ChIP-seq. Library preparation and initial bioinformatic analyses were completed by NOVOGENE. The construction of libraries involved the following steps. Four independent biological replicates were pooled per sample (5 OD600 INPUT samples, 15 OD600 for H3K9me2 ChIP IPs). ChIP DNA fragments were A-tailed before they were ligated with Illumina Compatible adapters. After size selection, libraries were amplified by PCR, pooled, and sequenced using the NovaSeq-6000. Read quality was assessed with FastQC (https://www.bioinformatics.babraham.ac.uk/projects/fastqc/). Adapter/index trimming was done using TRIMMOMATIC (*68*). Trimmed reads were mapped to the *S. pombe* reference genome (ASM294v2) with BWA (*69*) and a BAM file was generated. Normalization was done using bamCoverage (reads per kilobase million (RPKM)). Read counts per kilobase were adjusted by the length of the region and normalized to the total number of reads in the sample. This accounts for differences between samples in sequencing depth (*70*). Data was visualized using the Integrative Genomics Viewer (IGV) (*71–73*) where read amounts were separated in 50 bp.

**Small RNA (sRNA) sequencing**

Total RNA was extracted from the yeast cells using the hot phenol method as described (*74*), and the small RNA fraction purified from this was sequenced as follows (Fig.S3A). In short, 60 µg of total RNA was electrophoresed on a denaturing polyacrylamide gel (0.5×TBE, 17.5% acrylamide/bisacrylamide 19:1, 7 M urea) together with ZR small-RNA ladder (Zymo Research, R1090), the gel corresponding to 17 to 29 nt size of the marker was excised and RNA was extracted from the excised gel with ZR small-RNA PAGE recovery kit (Zymo Research, R1070).

The quality of the eluted RNA was evaluated by 5' end labelling with T4 Polynucleotide kinase (Thermo Fisher Scientific, EK0031) using its Buffer B conditions and 10 µCurie [$^{32}$P-γ]-ATP (Hartmann Analytic, SRP401) as per the recommendations of the kit. The labelled RNA was then resolved on a denaturing polyacrylamide gel as described above, electro-transferred on 0.45 µm Neutral Nylon Transfer Membrane (GVS, 1213403) using the Biorad MiniProtein II system, exposed to Imaging plate (Fujifilm, BAS-IP2025) and scanned on a TyphoonTM FLA9000 phosphor-images (GE Healthcare).

The eluted RNA were subsequently used for small RNA library preparation using small RNA-seq library Prep Kit (Lexogen GmbH, 052) as per the instructions of the manufacturer. The PCR products were analysed on a non-denaturing polyacrylamide gel (0.5×TBE, 17.5% acrylamide/bisacrylamide 19:1) to check the quality of the library and sequenced with Illumina NovaSeq X Plus system at Functional Genomic Center Zürich (FGCZ).

Demultiplexing was performed using the Illumina bcl2fasq Conversion software. Individual library sizes ranged from 43 to 100 million paired end 150 nt reads. Read quality was inspected using FastQC and MultiQC (*75*). Read 1 FASTQ files were trimmed and filtered using fastp (*76*), and aligned to the *S. pombe* reference genome (ASM294v2) using Bowtie 2 (*77*). BAM files were analyzed in R (version 4.5.0) using the Rsamtools (https://bioconductor.org/packages/release/bioc/html/Rsamtools.html) and GenomicAlignments (https://bioconductor.org/packages/release/bioc/html/GenomicAlignments.html) R Bioconductor packages.

To quantify the number of sRNA derived from centromeric repeats in Fig. S3D, the 20 centromere siRNA sequences used in Kawakami et al. (*44*) were applied.

**Statistical analyses of the partitioning patterns of the plasmids**

The question we were interested in answering in this study is whether DNA circle clustering causes its asymmetric partitioning. The null hypothesis of our statistical test is that the DNA circles do not cluster and their partitioning is random, specifically (i) upon cell division, each plasmid dot is partitioned into either daughter cells randomly, hence always with a probability of $p = 0.5$ and under no circumstances this probability changes; (ii) each partitioning outcome is independent of all the others. We did not distinguish the two daughter cells based on our observation (Fig.S1A-E). Given the premises of the experiments investigating the partitioning patterns of plasmids, we modelled the partitioning pattern under the null hypothesis and tested it against the measured data. We can consider each dot inheritance event as an independent Bernoulli trial with either success or failure. Suppose a cell before mitosis contains a dot number *d*. After mitosis, one of the daughter cells inherits a dot number *k*, and the other daughter inherits a dot number *d–k*. In this manner, the number of dots *d* prior to mitosis becomes the fixed number of Bernoulli trials, *k* becomes the number of successes, and *d–k* is the number of failures. Given these assumptions, the binomial distribution is the probability mass function that describes the pattern of partitioning.

For Fig.1F, we quantified the partitioning patterns of transformed cells containing zero to 20 dots and compared the experimental and theoretical frequencies of each category (fully asymmetric, $k > d-k = 0$; partially asymmetric, $k > 2 \times (d-k)$, $d \neq k$; symmetric, $k \leq 2 \times (d-k)$, $d \neq 0$, $k \neq 0$). For Fig.1G;3D;4F;5C;S6A, we calculated the probability mass function using the following formulas: $P_{k|d} = \frac{1}{2^{d-1}} \times \frac{d!}{k!(d-k)!}$, where $k = 0,\ldots,\frac{d-1}{2}$ (if *d* is odd) or $k = 0,\ldots,\frac{d}{2} - 1$ (if *d* is even); $P_{k|d} = \frac{1}{2^d} \times \frac{d!}{k!(d-k)!}$, where $k = \frac{d}{2}$ and *d* is even. Experimental frequency was calculated as follows: $F_{k|d} = \frac{n_{k|d}}{n_d}$, where $n_d$ is the number of cells observed with the number of dots *d*, and $n_{k|d}$ is the number of cells in the category *d* with *k* inherited dots. We restricted *d* from 1 to 20, except for the segregation pattern of RNAi mutants (Fig.3D), whereas *d* was restricted from 1 to 6. To obtain the probability of fully asymmetric inheritance $k_0$ dots in all *d*, we calculated the following equations: For theoretical probability, $P_{k_0 \cap d} = \sum_1^d P_{k|d} \times P_d = \sum_1^d \frac{1}{2^{d-1}} \times \frac{n_d}{n} = P_0$; for experimental probability, $F_{k_0 \cap d} = \sum_1^d F_{k|d} \times P_d = \sum_1^d \frac{n_{k_0|d}}{n_d} \times \frac{n_d}{n} = F_0$, where *n* is the total number of cells, $n_d$ is the number of cells observed with the number of dots *d*, $n_{k_0|d}$ is the number of cells with $k = 0$ inherited dots in the category *d*, and $n_{k_0}$ is the total measured number of cells with $k =$

0. We used the binomial test to investigate whether $F_0$ rejects $P_0$. In the binomial test, $P_0$ is the probability of success, $n_{k_0}$ is the number of successes, and *n* is the sample size (Bernoulli trials). To calculate the *p*-value, we calculate the following binomial complementary cumulative distribution function: *p*-value = $P(X \geq n_{k_0}) = 1 - P(X < n_{k_0}) = 1 - \sum_{i=0}^{n_{k_0}-1} P(X = i) = 1 - \sum_{i=0}^{n_{k_0}-1} \frac{n!}{i!(n-i)!} P_0^i (1 - P_0)^{n-i}$. We calculated the *Z*-score with the following equation: $Z = \frac{n_{k_0} - nP_0 + \frac{1}{2}}{\sqrt{nP_0(1-P_0)}}$.

**Figure S1**

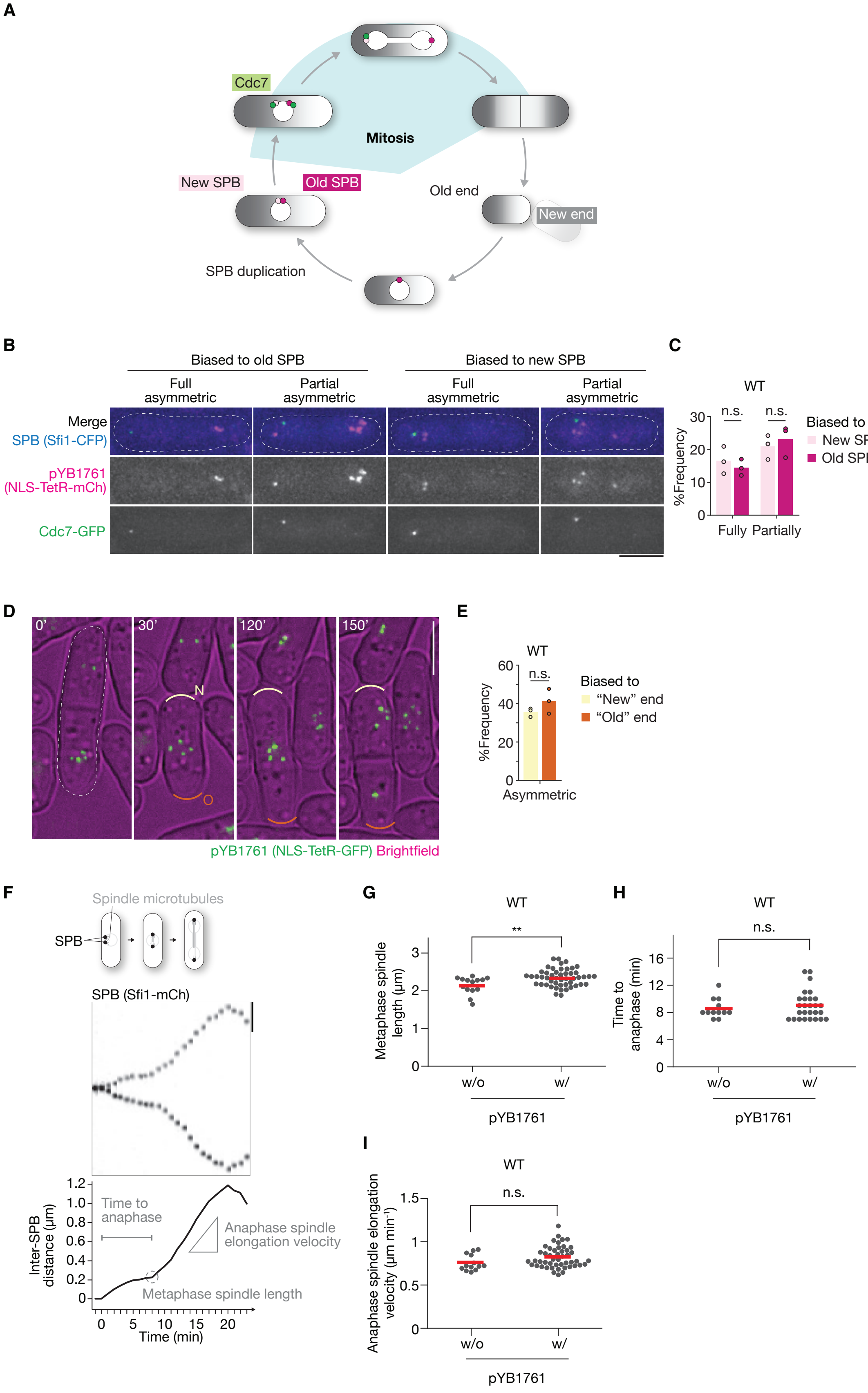


182

**Fig. S1. Unbiased asymmetric partitioning and effects on mitotic progression of the plasmid, related to Fig. 1.** (**A**) Cartoon of intrinsic structural biases in fission yeast: new and old SPBs, and new and old cell ends. The new SPB can be distinguished from the old SPB by the exclusive localization of Cdc7 on the new SPB in anaphase. (**B**) Microscopy images of cells showing each indicated partition pattern of pYB1761 (magenta) with SPB (blue) and Cdc7 (green). Scale bar, 5 µm. (**C**) Quantification of the fraction of each indicated partition pattern. $n > 166$ cells were tested in each trial. $n > 166$ cells were tested in each trial. $N = 3$ trials. n.s., not significant (Student's *t*-test). (**D**) Microscopy images of a cell showing asymmetric partitioning of pYB1761 (green) towards the new cell end. The new and old cell ends are depicted as 'N' and 'O' at 30 min, respectively. The white dotted line in 0 min represents the outline of the cell focused on the following time points. Scale bar, 5 µm. (**E**) Quantification of the fraction of each indicated partition pattern. $n > 106$ cells were tested in each trial. $N = 3$ trials. n.s., not significant (Student's *t*-test). (**F**) Upper, localization of SPB was observed every 1 min throughout mitosis in the wild-type cell transformed with pYB1761. The trajectory of two SPBs is shown as a kymograph. Scale bar, 2 µm. Lower, a representative plot of the kinetics of the inter-SPB distance of the upper image is shown. Time point 0 min corresponds to the onset of mitosis. Metaphase spindle length is defined as the length of the inter-SPB distance just before the two SPBs resume separation. Time to anaphase is defined as the duration between the time point just before the two SPBs start separation and the time point just before they resume separation. Anaphase spindle elongation velocity is defined as the slope from the time point at which two SPBs resume separation to the time point that the inter-SPB distance reaches its maximum in the kinetics of inter-SPB distance. (**G**) Quantification of metaphase spindle length measured as the inter-SPB distance in wild-type cells with and without pYB1761. $n = 63$ cells. The red lines represent the mean value. $^{**}p < 0.01$ (Student's *t*-test). (**H**) Quantification of time from mitotic onset to anaphase onset in wild-type cells with and without pYB1761. $n = 39$ cells. The red lines represent the mean value. n.s., not significant (Student's *t*-test). (**I**) Quantification of the velocity of anaphase spindle elongation measured in wild-type cells with and without pYB1761. $n = 61$ cells. The red lines represent the mean value. n.s., not significant (Student's *t*-test).

**Figure S2**

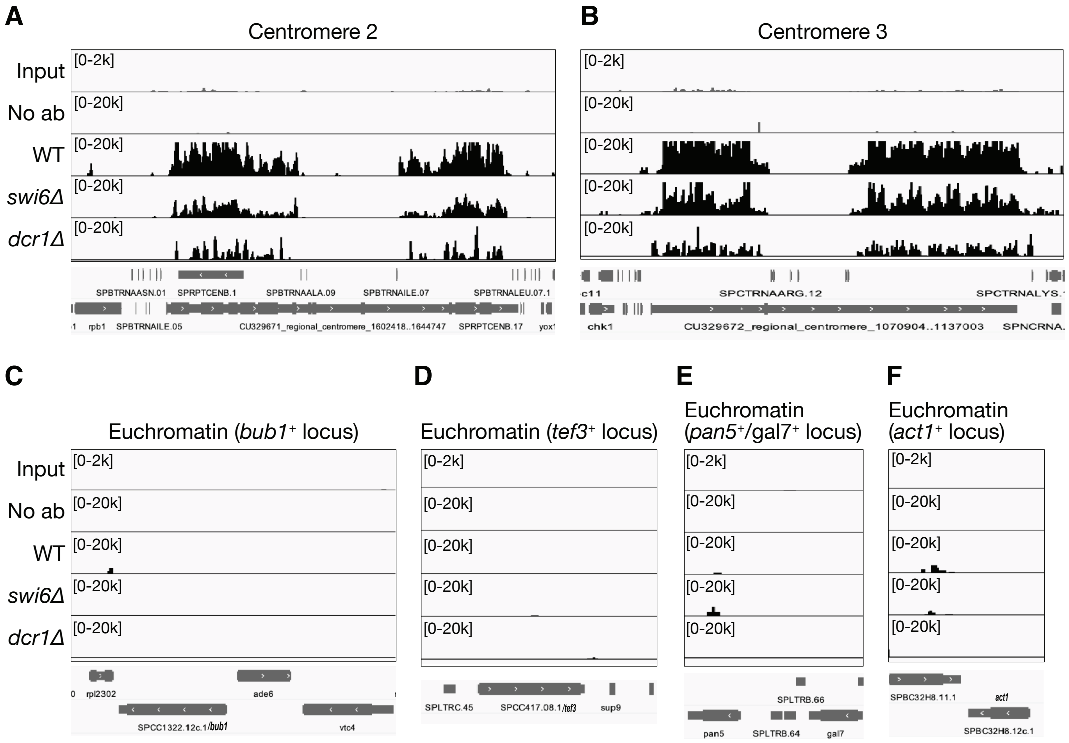


**Fig. S2. Profiles of H3K9me2 ChIP-seq on chromosomal regions, related to Fig. 2.** Profiles on H3K9me2 ChIP-seq on (A) the chromosome 2 centromere, (B) the chromosome 3 centromere, and (C-F) euchromatin regions for the indicated genotypes. No ab, negative control without antibodies.

**Figure S3**

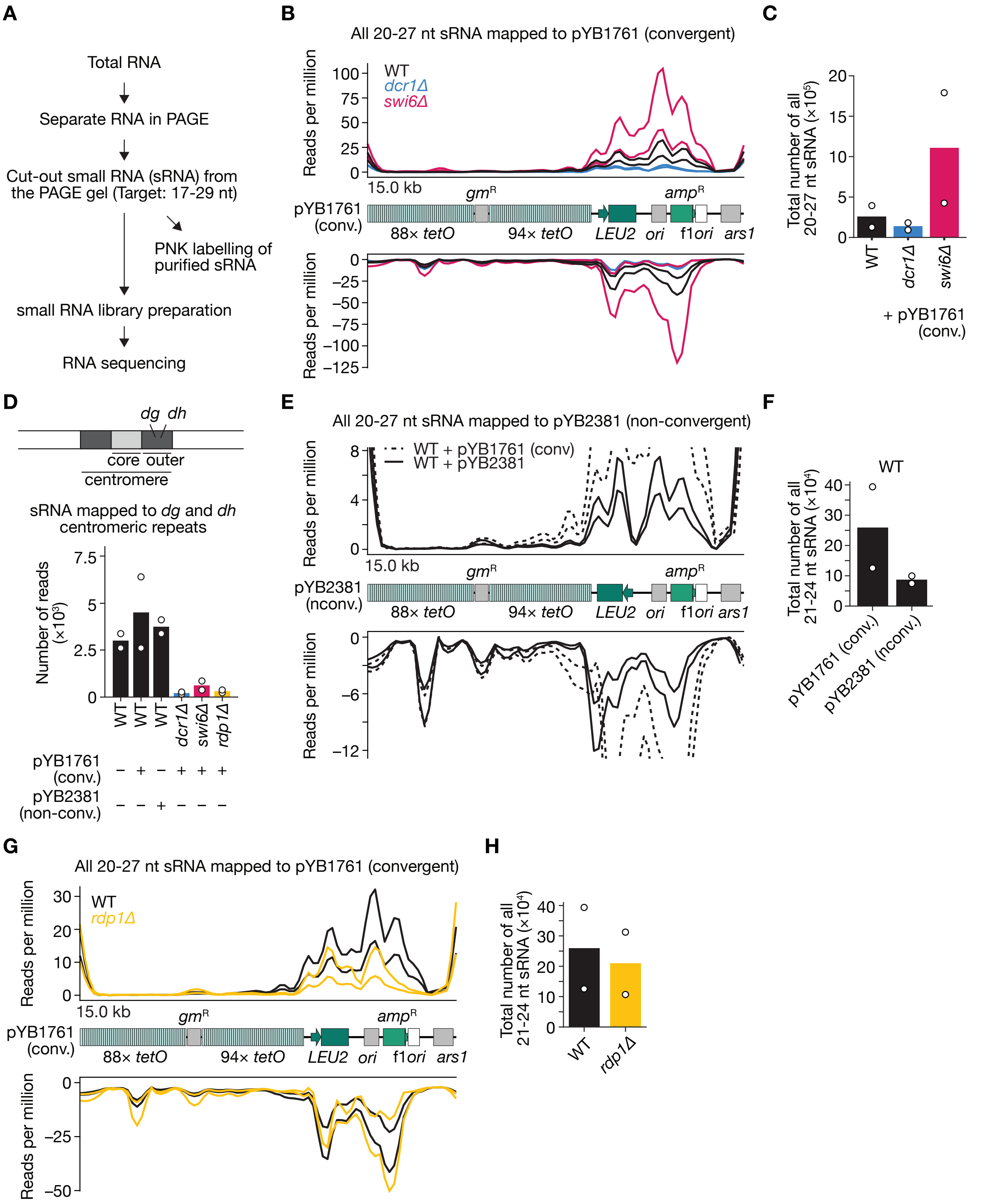


**Fig. S3. Profiles of non-biased 20-27 nt sRNA mapped to plasmids, related to Fig. 2 and 4.** (**A**) Flow chart of sample preparation for sRNA sequencing. (**B**) Profile of all 20-27 nt sRNA from the indicated genotype mapped to pYB1761. *N* = 2 trials for each of the indicated genotypes. (**C**) Plot of the total number of all 20-27 nt sRNA mapped to pYB1761 in each genotype. *N* = 2 trials for each of the indicated genotypes. (**D**) Plot of the total number of sRNA mapped to *dg* and *dh* centromeric repeats in each condition. *N* = 2 trials for each of the indicated genotypes. For the sRNA sequences, see Materials and Methods. (**E**) Profile of all 20-27 nt sRNA from wild-type cells mapped to pYB1761 (dotted lines) or pYB2381 (solid lines). *N* = 2 trials for each condition. (**F**) Plot of the total number of all 20-27 nt sRNA mapped to pYB1761 or pYB2381 in wild-type cells. *N* = 2 trials for each of the indicated genotypes. (**G**) Profile of all 20-27 nt sRNA from the indicated genotype mapped to pYB1761. *N* = 2 trials for each of the indicated genotypes. (**H**) Plot of the total number of all 20-27 nt sRNA mapped to pYB1761 in each genotype. *N* = 2 trials for each of the indicated genotypes.

**Figure S4**

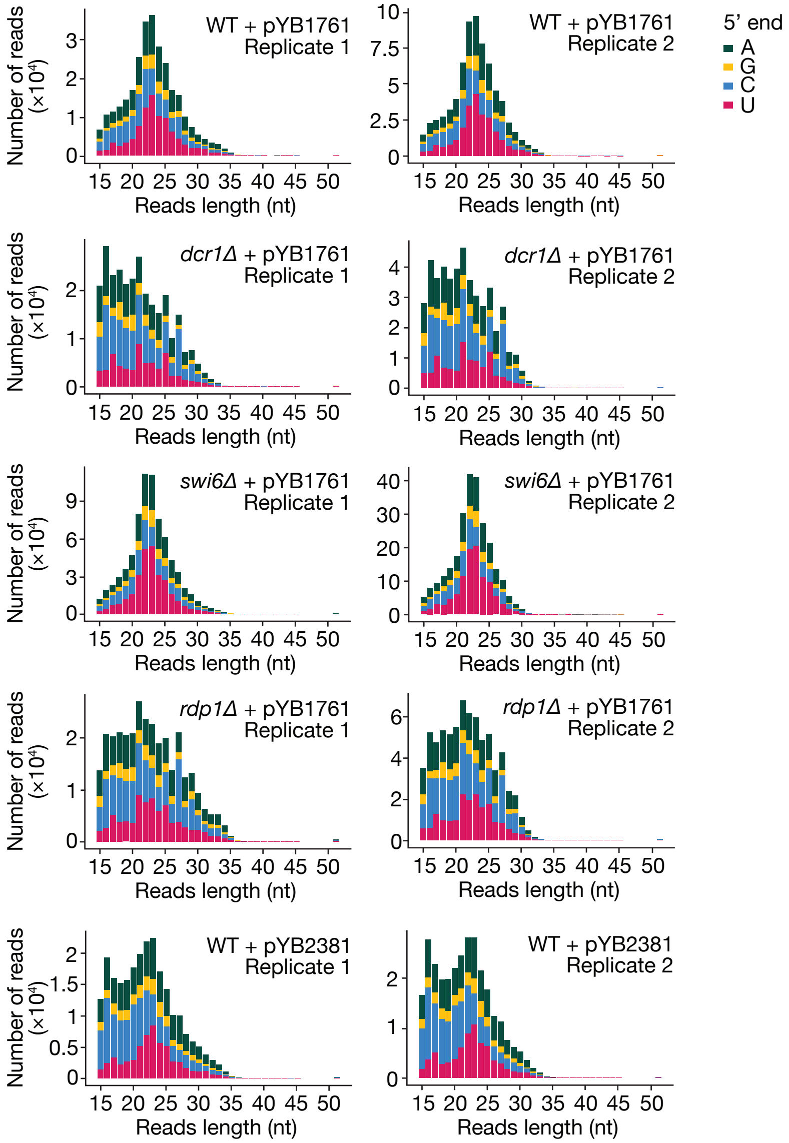


**Fig. S4. 5'-terminal nucleotide composition of deep-sequenced sRNAs of all biological replicates.**

**Figure S5**

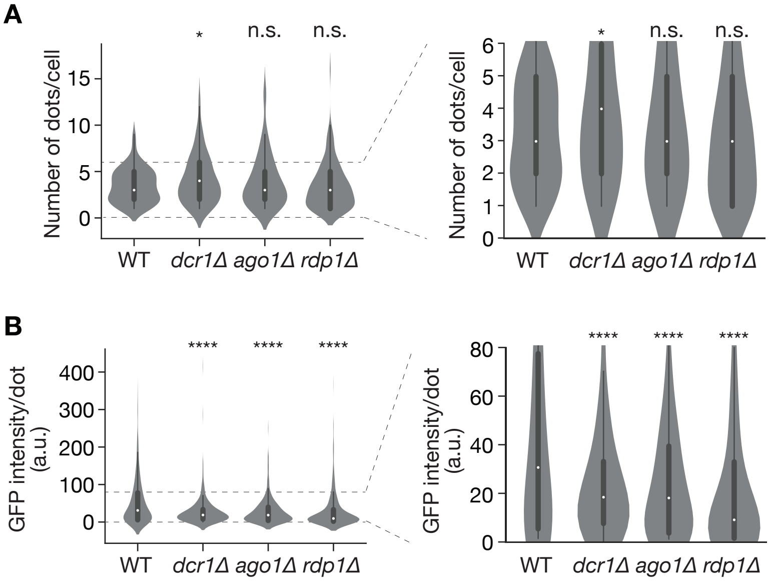


**Fig. S5. Clustering of pYB1761 in RNAi mutants, related to Fig. 3.** (**A**) Left, quantification of the number of fluorescent dots in the indicated genotypes. Right, same as the left, except zoomed in on the indicated range. $n > 92$ cells. A white circle in each violin plot represents the mean value. n.s. not significant; $^{*}p < 0.05$ (Welch's t-test). (**B**) Left, quantification of fluorescent intensity of each dot in the indicated genotypes. Right, same as the left, except zoomed in on the indicated range. $n > 319$ dots were tested. A white circle in each violin plot represents the mean value. $^{****}p < 0.0001$ (Welch's *t*-test).

**Figure S6**

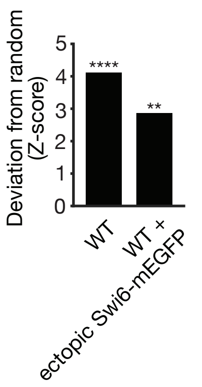


**Fig. S6. Asymmetric partitioning of pYB1761 in cells expressing Swi6-mEGFP ectopically, related to Fig. 5.** Quantification of the divergence of the distribution of pYB1761 partition patterns from the expected in indicated cells. Data for wild-type cells is the same as Fig. 1G. $n$ = 177 cells for ectopic Swi6-mEGFP. $^{**}p < 0.01$; $^{****}p < 0.0001$ (binomial test analyses). For details, see Materials and Methods.

**Figure S7**

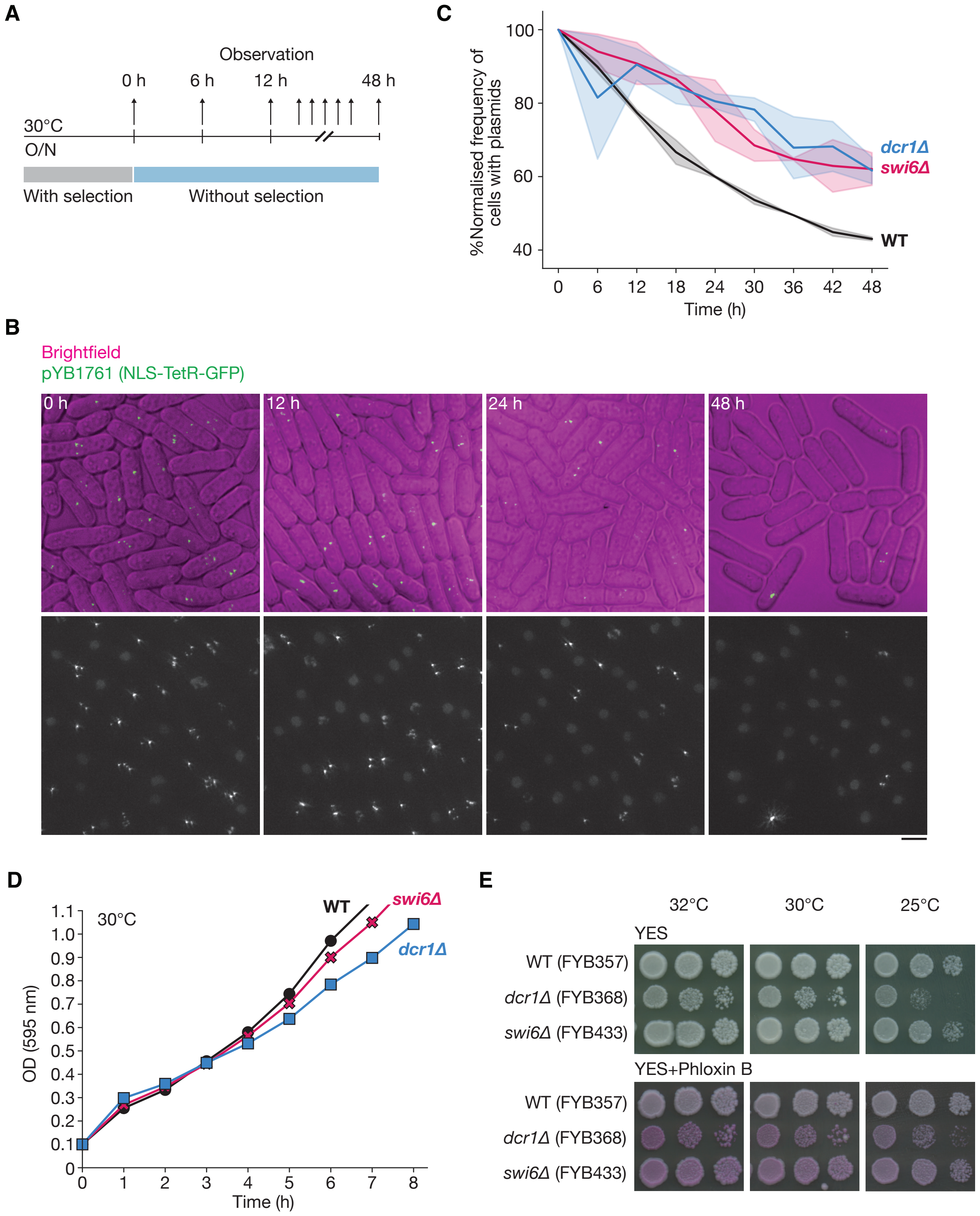


**Fig. S7. Heterochromatinization of the plasmid promotes its elimination from the cell population.** (**A**) Schematic overview of experiments. (**B**) Microscopy images of wild-type cells with pYB1761 (green) with brightfield (magenta) at the indicated time points. Scale bar, 5 μm. (**C**) Quantification of the fraction of cells retaining plasmids at each time point. Lines and shaded areas represent the mean and 95% confidence interval from two independent experiments, respectively. *n* > 225 cells were tested at each time point in the indicated genotype. (**D**) OD (595 nm) measurement of log-phase cultures at 30°C. (**E**) Ten-fold serial dilutions of the indicated strains were spotted on YES and YES including 2 μg/mL of Phloxin B, and incubated at the indicated temperature for 3 days.

### Figure S8

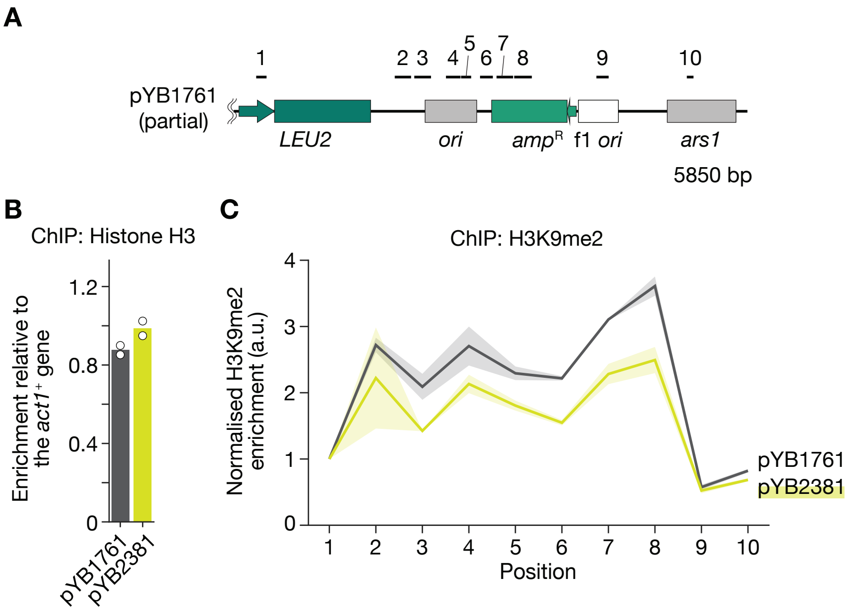


**Fig. S8. Impacts of plasmid-intrinsic transcription features on heterochromatinization of the plasmid.** (**A**) Schematic overview of targeted regions by ChIP-qPCR. (**B**) Histone H3 on the *LEU2* promoter ("1" in Fig.S8A) of pYB1761 and pYB2381 was detected by ChIP-qPCR in log-phase wild-type cells. $N = 2$ experiments. (**C**) H3K9me2 on the indicated regions of pYB1761 and pYB2381 was detected by ChIP-qPCR in log-phase wild-type cells. Lines and shaded areas represent the mean and 95% confidence interval from two independent experiments, respectively. Enrichment values of targeted regions relative to euchromatin genes (see Methods) were normalized by the value of the "1" region.

**Figure S9**

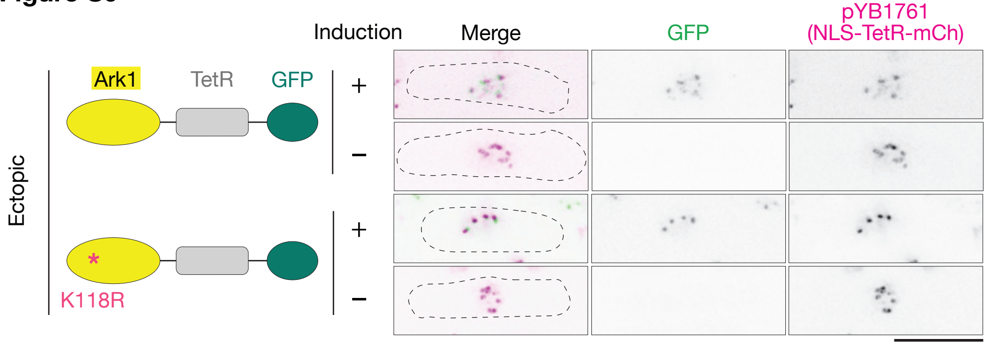


**Fig. S9. Microscopy images of the Ark1 tethering experiments, related to Fig. 5.** Microscopy images of pYB1761 (magenta) and ectopically expressed Ark1 fusion proteins (green) in cells. Scale bar, 5 µm.

**Table S1.**

S. pombe strains used in this study. Nomenclatures are based on *(57)*.
Table S1 is provided as a separate file (TableS1_Ebina_et_al_bioRxiv.xlsx).

**Table S2.**

Plasmids used in this study.

| Name | Purpose | Source |
|---|---|---|
| pYB1678 | Inserting *P81nmt1-NLS-tetR-GFP* at the *ura4* locus | This study |
| pYB1761 | Convergent plasmid | This study |
| pYB2114 | Inserting *P.nup211-NLS-tetR-mCherry* at the *ura4* locus | This study |
| pYB2125 | Inserting *P.nup211-NLS-tetR-GFP* at the *ura4* locus | This study |
| pYB2381 | Non-convergent plasmid (the direction of the *LEU2* is flipped from pYB1761) | This study |
| pYB4042 | Inserting *P41nmt1-ark1-NLS-tetR-GFP:bsdR* at the *z2* locus on chromosome II | This study |
| pYB4084 | Inserting *P41nmt1-ark1(K118R)-NLS-tetR-GFP:bsdR* at the *z2* locus on chromosome II | This study |
| pYB4084 | Inserting *P41nmt1-ark1-NLS-GFP:bsdR* at the *z2* locus on chromosome II | This study |
| pYB4098 | Template of PCR amplification of the DNA fragment coding *kanMX6:P81nmt1-swi6-(SG4)3S-mEGFP:T.ADH1* | This study |

**Table S3.**

Primers used in this study.

Table S3 is provided as a separate file (TableS3_Ebina_et_al_bioRxiv.xlsx).
